# Increased intracellular crowding during hyperosmotic stress

**DOI:** 10.1101/2022.12.30.522363

**Authors:** Akira Kitamura, Sho Oasa, Haruka Kawaguchi, Misato Osaka, Vladana Vukojević, Masataka Kinjo

## Abstract

Hyperosmotic stress activates in live cells numerous processes and also promotes intracellular protein/RNA aggregation and phase separation. However, the time course and the extent of these changes remain largely uncharacterized. To investigate dynamic changes in intracellular macromolecular crowding (MMC) induced by hyperosmotic stress in live cells, we used Fluorescence Lifetime Imaging Microscopy (FLIM) and Fluorescence Correlation Spectroscopy (FCS) to quantify changes in the local environment by measuring the fluorescence lifetime and the diffusion of the monomeric enhanced Green Fluorescent Protein (eGFP), respectively. Real-time monitoring of eGFP fluorescence lifetime showed that a faster response to environmental changes due to MMC is observed than when measuring the acceptor/donor emission ratio using the MMC-sensitive Förster Resonance Energy Transfer sensor (GimRET). This suggests that eGFP molecular electronic states and/or collision frequency are affected by changes in the immediate surroundings due to MMC without requiring conformational changes as is the case for the GimRET sensor. Furthermore, eGFP diffusion assessed by FCS indicated higher intracellular viscosity due to increased MMC during hyperosmotic stress. Our findings reveal that changes in eGFP fluorescence lifetime and diffusion are early indicators of elevated intracellular MMC. These variables can therefore be used for quantitative characterization of MMC in live cells.

## Introduction

Cells dynamically interact with their surroundings and changes in the extracellular environment are readily accompanied by adjustments of intracellular properties through the activation of numerous biochemical reaction pathways. For example, changes in solutes concentration outside the cell lead to altered movement of water across the plasma membrane, disrupting the intracellular osmolarity that is tightly regulated by balancing the intake and excretion of water and electrolytes^1,2^. If an increase in solutes concentration is observed, hyperosmotic stress ensues, which, if not interfered with, can rapidly give rise to protein damage due to cellular water loss^3^. Cells activate various biochemical pathways in a non-steady-state manner to counteract the hyperosmotic stress^2^. Post exposure to a hyperosmotic environment, cells shrink due to dehydration, *i*.*e*., due to the loss of water molecules from the cell to the surrounding. To diminish or shut down unwanted and unnecessary nascent polypeptide synthesis, aggregation of macromolecules leads to the formation of a dense network of interactions due to dehydration, strong enough to drive phase separation, resulting in the formation of stress granules (SGs) containing mRNA, RNA-binding proteins (RBPs) and ribosomes^4-6^.

During hyperosmotic stress the cell volume changes to such an extent that intracellular macromolecular crowding (MMC) is considered to also change. To read out the MMC status at different locations in the cells, the Förster resonance energy transfer (FRET)-based sensor GimRET, a fusion protein between the cyan fluorescent protein (CFP) and the crowding-sensitive mutant of the yellow fluorescent protein (YFP) with an inserted glycine residue before Tyr145, was recently designed^7^. The aim of this work is twofold: to use GimRET to characterize MMC during hyperosmotic stress and to establish an alternative, fluorescence lifetime-based method for *in situ* monitoring of MMC changes during hyperosmotic stress in real-time.

Fluorescence lifetime (τ_FL_), *i*.*e*., the lifetime of a molecule in the excited singlet electronic state, is an intrinsic property of the fluorophore that does not depend on fluorophore concentration, excitation, and/or photobleaching, but rather depends on environmental factors, such as temperature, solvent polarity, and the presence of molecules with the capacity to quench fluorescence. It is therefore well suited to study environmental changes, such as MMC in live cells. τ_FL_ is defined as the mean time required for a population of molecules in the excited state to be reduced to 36.8 % of its original value (*i*.*e*. by a factor of Napier’s constant, *e* = 2.718), by losing energy through fluorescence emission, which is a radiative process, and by various nonradiative processes, such as intersystem crossing to the triplet state, internal conversion, collisional, *i*.*e*. dynamic quenching, energy transfer, charge transfer, *etc*.^8,9^ As such, τ_FL_ is reciprocally proportional to the total decay rate, comprising contributions from the radiative and all non-radiative rate constants. Theoretically, however, FRET between identical fluorophores (homo-FRET) does not affect the fluorescence lifetime of the probes.^10^ Hence, τ_FL_ decreases when the rate constant of any of the non-radiative processes increases. τ_FL_ can be measured in either the frequency or the time domain. Here, time domain measurements were used^11,12^ to probe the local cellular environment^13^; FRET-donor and -acceptor ratiometric imaging using the GimRET sensor was deployed to monitor the changes in MMC after cell exposure to hyperosmotic stress; and diffusion of eGFP was measured using fluorescence correlation spectroscopy (FCS)^6,14,15^, to acquire information about local viscosity^16^.

## Results

### Fluorescence lifetime of eGFP rapidly decreased and slowly recovered in cells exposed to hyperosmotic stress

CLSM imaging revealed that eGFP transiently expressed in murine neuroblastoma Neuro-2a (N2a) cells is uniformly distributed inside the cells (Figure 1A, upper row). Following the exposure to hyperosmotic stress caused by 290 mM NaCl (Figure 1A, lower row), the morphology of the N2a cells markedly changed within some 20 s, as evident from the transmission bright field images in Figure 1A. As before, eGFP was uniformly localized in both, the cytoplasm and the cell nucleus, however, in a significant portion of N2a cells a pronounced eGFP fluorescence rim was observed at the boundary between the nucleus and the cytoplasm (Figure 1A, lower row and Supplemental Figure S1, lower row, white arrows). In addition, formation of SGs was observed in the cytoplasm, verified using the SG marker T cell intracellular antigen 1 tagged at the N-terminus with the Red Fluorescent Protein variant mCherry (mCherry-TIA1; Figure 1A).

**Figure 1.**
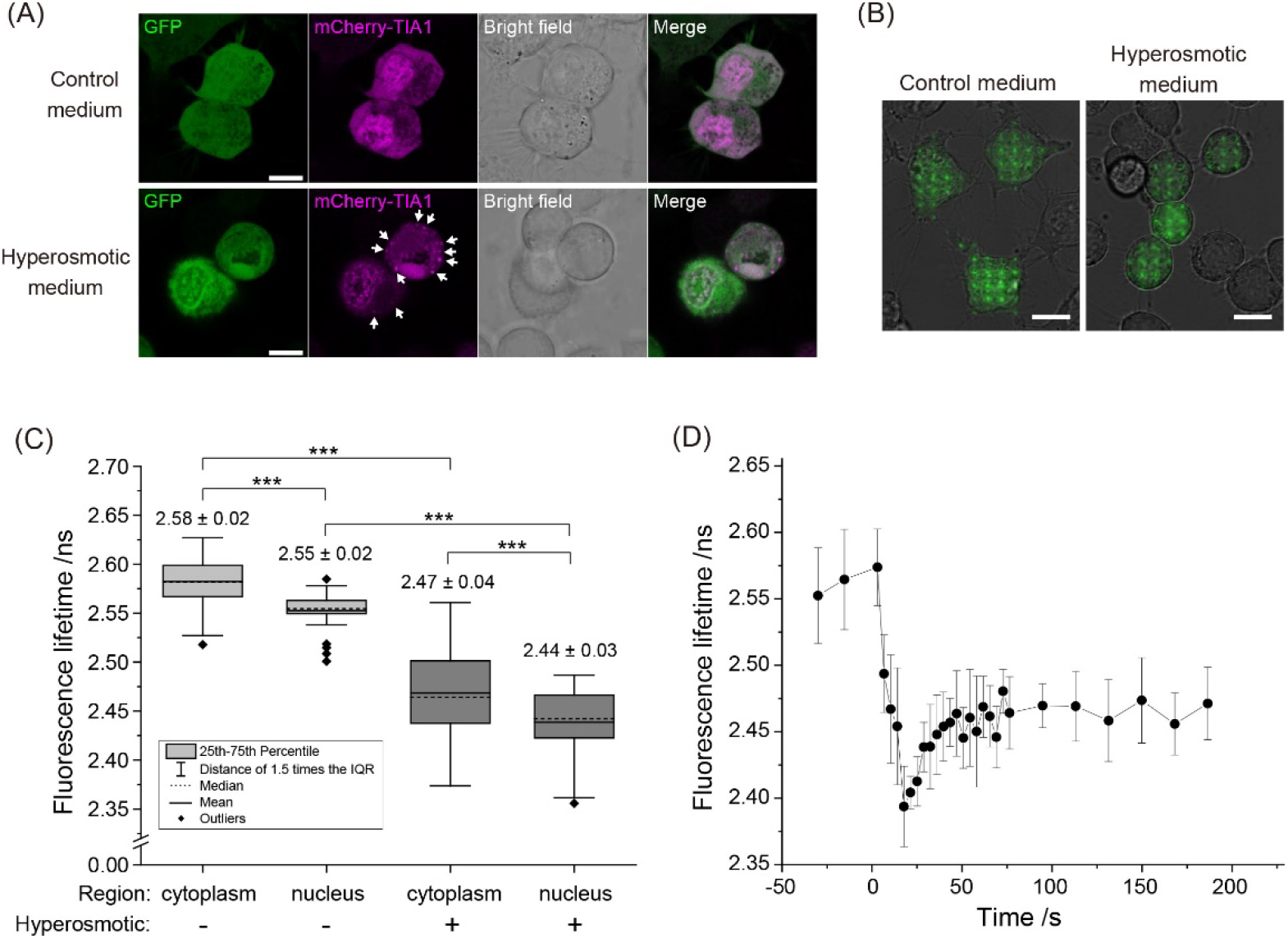
Hyperosmotic stress-induced changes in macromolecular crowding (MMC) followed by phase separation observed through the formation of stress granules (SGs) and environmental changes reflected by variations in eGFP fluorescence lifetime. **(A)** Fluorescence and transmitted light images of live N2a cells transiently expressing eGFP monomers and TIA1-mCherry when grown under normal conditions (upper row) and upon exposure to hyperosmotic stress (lower row). The images were acquired using Confocal Laser Scanning Microscopy. Scale bar: 10 μm. The white arrows point to stress granules (SGs). **(B)** Fluorescence images of eGFP-expressing N2a cells acquired using the home-built microscope for massively parallel fluorescence lifetime imaging (mpFLIM) superimposed on corresponding brightfield images. Scale bar: 10 μm. Bright green spots indicate individual excitation foci where FLIM measurements were simultaneously performed. **(C)** Box and whisker plots showing eGFP fluorescence lifetime in the cytoplasm and the nucleus of N2a cells under normal conditions (light grey) and during hyperosmotic stress (dark grey). The numbers on the box graph show mean ± SD. The number of independent spots in which eGFP fluorescence lifetime was measured were: 87 in the cytoplasm and 37 in the nuclei of 8 cells under normal physiology, and 84 spots in the cytoplasm and 70 in the nuclei of 15 cells exposed to hyperosmotic stress. Statistical significance was tested by Student’s *t* test: \*\*\**p* < 0.001. **(D)** Time course of changes in eGFP fluorescence lifetime in live N2a cells before and after exposure to hyperosmotic stress, recorded using the mpFLIM system. The hyperosmotic stress medium was added at 0 s. Dots and bars show mean ± SD (the number of analyzed spots from 5 cells, n = 9).

We measured eGFP fluorescence lifetime (τ_FL,eGFP_) in live N2a cells under normal growth conditions (Figure 1B, left) and hyperosmotic stress (Figure 1B, right) using a home-built massively parallel FLIM system (mpFLIM) without laser scanning in which 16×16 confocal foci were generated using a Diffractive Optical Element (DOE) to split the incident laser beam into a 2-dimensional (2D) array of beams and a 2D Single Photon Avalanche Diodes (SPAD) array camera with matching geometry^12^. Subcellular positions where FLIM measurements were performed were identified using bright field and fluorescence imaging (Figure 1B). Under normal conditions, τ_FL,eGFP_ in the cytoplasm was 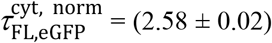 ns (mean ± SD), whereas a slightly shorter fluorescence lifetime was measured in the cell nucleus, 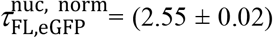 ns, (Figure 1C). Upon hyperosmotic stress, τ_FL,eGFP_ in both the cytoplasm and the nucleus markedly decreased, becoming significantly shorter, 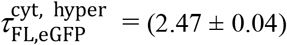 ns and 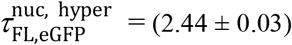 ns (Figure 1C). Moreover, τ_FL,eGFP_ in N2a cells swiftly decreased just after addition of the hyperosmotic medium and gradually recovered thereafter (Figure 1D, reproduced in Supplemental Figure S2A (black squares)), while τ_FL,eGFP_ in live N2a cells not exposed to hyperosmotic stress did not significantly change during the measurement (Supplemental Figure S2A (grey dots)). Concomitantly, fluorescence intensity, as reflected by photon counts, swiftly increased ∽ 1.45-times just after addition of the hyperosmotic medium and gradually decreased thereafter (Supplemental Figure S2B (black squares)). This slow decrease in fluorescence intensity that is due to eGFP photobleaching during signal acquisition for fluorescence lifetime measurements could also be observed in measurements on control cells, not exposed to hyperosmtic stress (Supplemental Figure S2B (magenta squares)). Of note, while good correlation between the swift increase in fluorescence intensity, as reflected by photon counts measurement (Supplemental Figure S2C (magenta triangles)), and the swift decrease in eGFP fluorescence lifetime (Supplemental Figure S2C (grey triangles)) was observed immediately after the addition of the hyperosmotic medium (*t* = 0 s), the ensuing slow-changing fluorescence intensity decrease/τ_FL,eGFP_ increase appeared to proceed at different paces – the maximum in fluorescence intensity was reached at the same time as τ_FL,eGFP_ minimum, but while the fluorescence intensity appeared to have settled (continuously decaying due to photobleaching), the τ_FL,eGFP_ continued to increase for some 50 s before reaching a plateau value (Supplemental Figure S2C). This suggests that while equilibration of eGFP concentration by diffusion is quickly achieved following hyperosmotic stress induction, slow intracellular adaptation processes continue to proceed, as reflected by τ_FL,eGFP_.

To examine whether the change in τ_FL,eGFP_ is primarily due to MMC or due to change in viscosity, control experiments were performed using eGFP in a series of solutions of bovine serum albumin (BSA), commonly used as a crowding agent in biochemical reactions^17^, with concentrations of BSA varying from 0–250 mg/ml (Supplemental Figure S3A), or with a different NaCl concentration (Supplemental Figure S4). We have observed that τ_FL,eGFP_ became shorter in a BSA-concentration-dependent manner (Supplemental Figure S3A), which is indicative of dynamic quenching of eGFP fluorescence in BSA solutions of increasing concentration, *i*.*e*. increasing MMC. To further test this interpretation, we have used the relationship 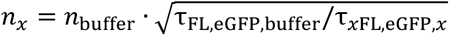 that is derived based on the Strickler–Berg equation^18^, to calculate the refractive index change based on changes in eGFP fluorescence lifetime (Supplemental Figure S3B), and compared these values with refractometry data from the literature where refractive index dependence on BSA concentrations was characterized^19^. A very good agreement between literature data and our *in vitro* analysis of BSA effect on τ_FL,eGFP_ is obtained (Supplemental Figure S3B, cyan *vs* orange dots). Moreover, our *in vitro* data on NaCl effect on τ_FL,eGFP_, showed that τ_FL,eGFP_ remains virtually unaltered at 290 mM NaCl, the concentration that was used in this study (Supplemental Figure S4). This is in line with reported refractive index measurements in NaCl solutions of different concentrations (Supplemental Figure S3B, green dots)^20^, based on which the expected decrease in τ_FL,eGFP_ using the *n_x_* is 0.01, which is within the experimental error of our measurements (SD value in Supplemental Figure S4).

### Y/C ratio of the MMC-sensitive GimRET sensor rapidly decreased and slowly recovered in cells exposed to hyperosmotic stress

To further assess whether MMC increases in cells during hyperosmotic stress, we monitored changes in MMC in live cells using the FRET-based MMC sensor GimRET^7^. In the GimRET sensor, quenching of YFP decreases FRET efficiency between CFP and YFP, resulting in a decrease in fluorescence intensity ratio between the YFP acceptor and the CFP donor (Y/C ratio) as MMC increases (Figure 2A). The subcellular localization pattern of GimRET (Figure 2B) was the same as for eGFP (Figure 1A) in both, the non-stressed (control) and stressed cells. In line with this, we have observed that the Y/C ratio of the GimRET sensor in live N2a cells abruptly decreased shortly after exposing the cells to the hyperosmotic medium, and then gradually recovered (Figure 2C & D). In contrast, the Y/C ratio of GimRET in live N2a cells not exposed to hyperosmotic stress was not changing during the timelapse measurement (Supplemental Figure S5).

**Figure 2.**
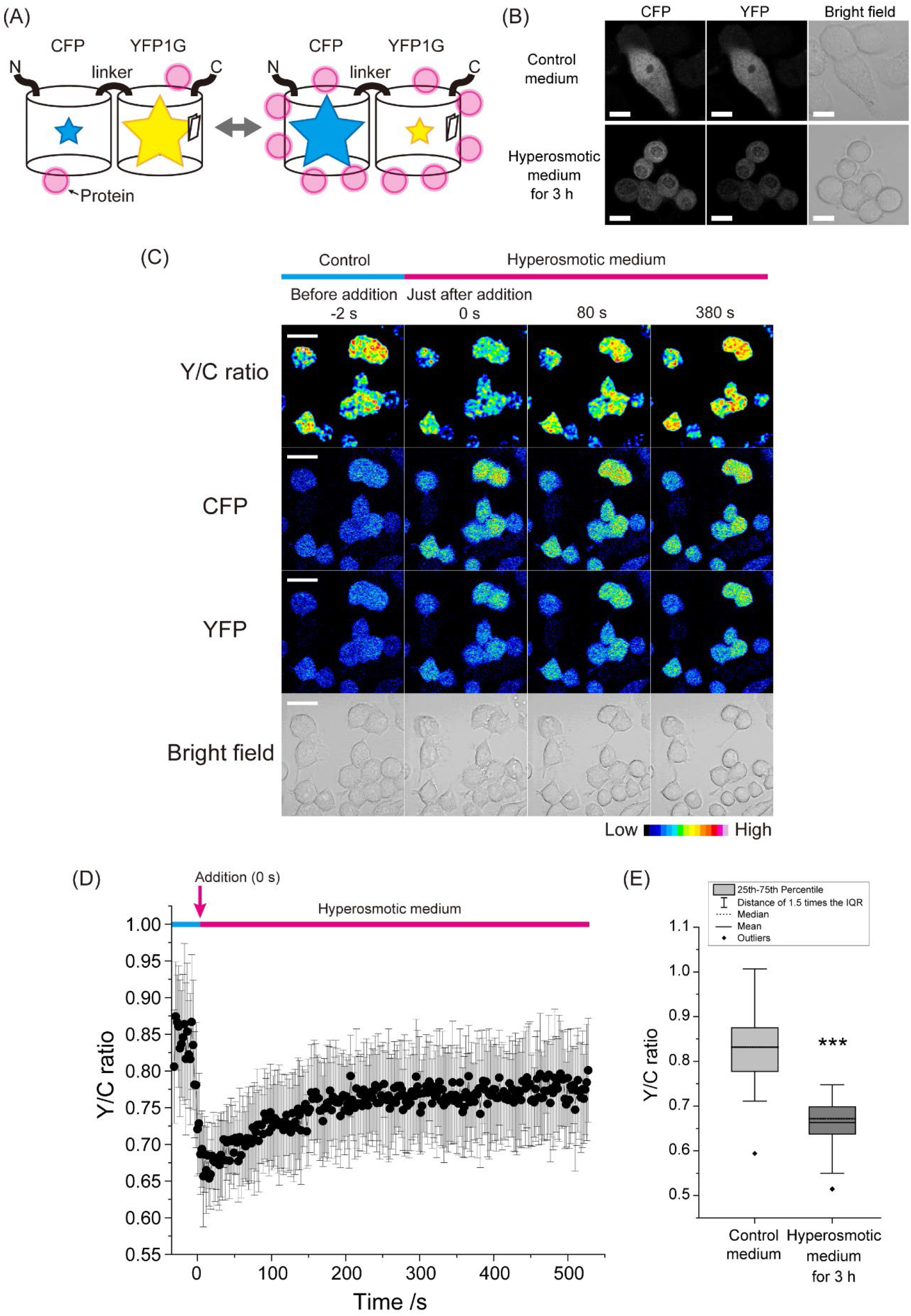
Hyperosmotic stress-induced changes in intracellular macromolecular crowding (MMC) monitored using the using the MMC-sensitive GimRET sensor. **(A)** Schematic drawing of the FRET-based MMC-sensitive sensor GimRET, a fusion protein between CFP and the crowding-sensitive mutant of YFP with an inserted glycine residue (YFP1G). The size of the star indicates relative fluorescence intensity under low/high MMC (magenta circles). **(B)** Confocal microscopy images of CFP and YFP fluorescence and transmission microscopy images of live N2a cells expressing the MMC-sensitive sensor GimRET when grown in regular cell culture medium (control) and after 3 hours in hyperopsmotic medium containing 290 mM NaCl. Scale bar = 10 μm. **(C)** Fluorescence time-lapse images before (-2 s) and after the addition of 290 mM NaCl to N2a cells expressing the MMC-sensitive sensor GimRET. Time of NaCl addition was at 0 s. The Y/C ratio was determined by dividing the background-corrected fluorescence intensity in the YFP channel by the background-corrected fluorescence intensity in the CFP channel and smoothed using the Gaussian blur function. The pseudo color scale is shown in the right bottom corner. Scale bar: 20 μm. **(D)** Corresponding changes in the Y/C ratio over time. Dots and bars indicate mean ± SD (the number of analyzed cells, n = 18). **(E)** Box and whisker plot showing Y/C ratio values measured in live N2a cells expressing the GimRET sensor after 3 hours incubation in hyperosmotic or control medium (the number of analyzed cells, n = 53). Statistical significance was tested by Student’s *t* test: \*\*\**p* < 0.001.

After incubation of the N2a cells in the hyperosmotic medium for 3 h, the Y/C ratio of GimRET was still lower than in the normal growth medium (Figure 2E). Cell shrinkage and morphology changes were also observed shortly after addition of the hyperosmotic medium, and remained visible throughout the experiment time as evident from the bright field image (Figure 2C, bottom row), showing cell volume recovery. Of note, the half recovery time of τ_FL,eGFP_ and Y/C ratio of GimRET after addition of stress medium was different, 9.7 s and 73.3 s, respectively (Figure 3).

**Figure 3.**
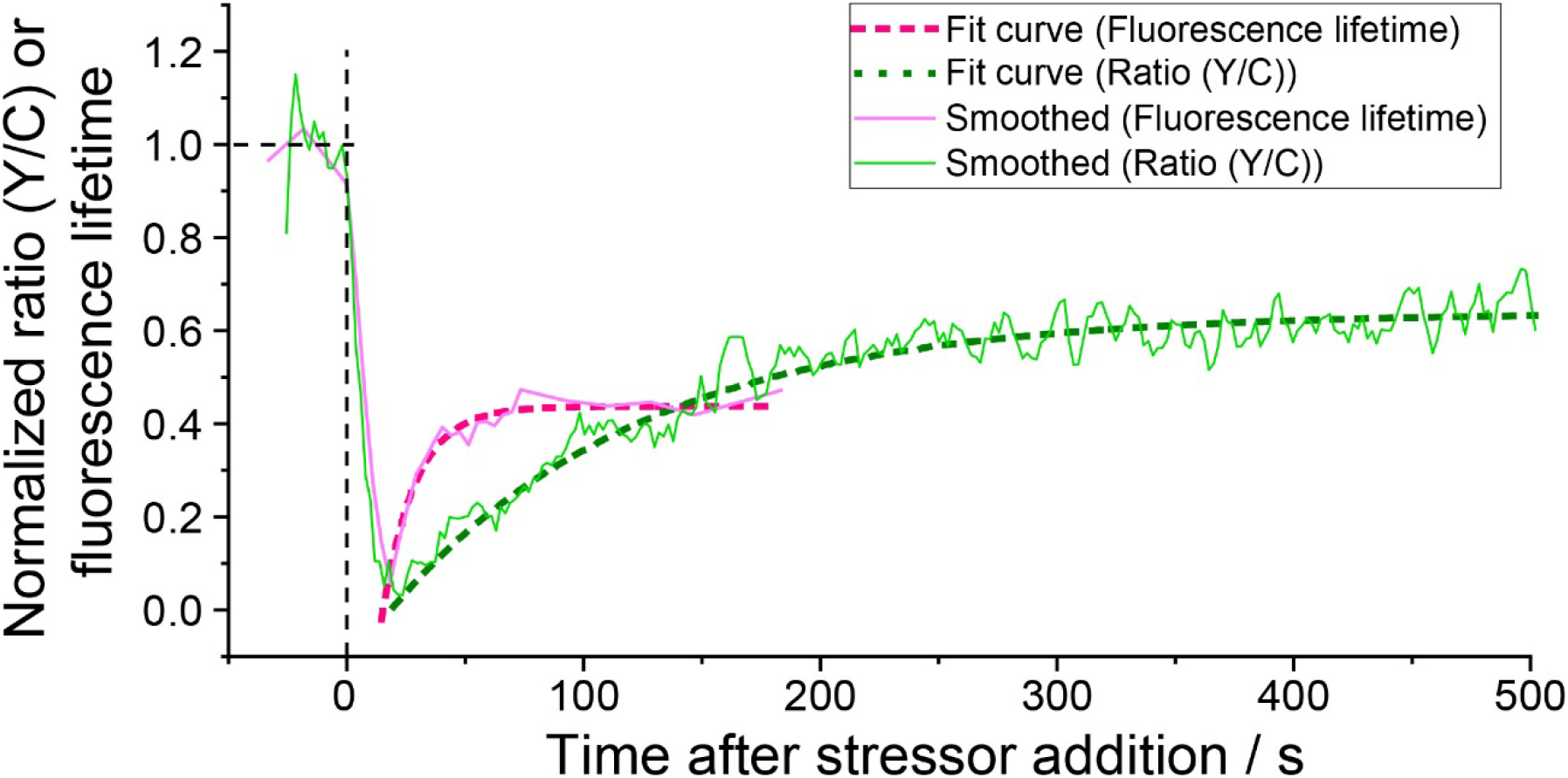
eGFP fluorescence lifetime is an earlier marker of hyperosmotic stress-induced macromolecular crowding (MMC) than the Y/C ratio of the MMC-sensitive GimRET probe. Normalized recovery curves showing changes in eGFP fluorescence lifetime (pink) and Y/C ratio of GimRET (green) caused by hyperosmotic stress-induced MMC. The normalized curves are derived from the data shown in Figure 1D and Figure 2B. The thin solid lines are smoothed using adjacent averaging of two data points The dashed lines show an exponential recovery model fitted to all data points.

### eGFP diffusion and anomalous diffusion parameter (α) decrease in cells exposed to hyperosmotic stress indicating an increase in intracellular viscosity and heterogeneity

To elucidate how MMC induced by hyperosmotic stress affects intracellular viscosity, the diffusion of monomeric eGFP was measured using FCS. We have observed that eGFP diffusion time, that is the decay time of the autocorrelation curve, was longer both in the cytoplasm and in the nucleus when measured in N2a cells exposed to hyperosmotic stress compared to N2a cells under normal physiology (Figure 4A). This indicates that eGFP diffusion is slower in N2a cells exposed to hyperosmotic stress. By fitting the experimentally derived autocorrelation curves using the one-component anomalous diffusion model, eGFP diffusion coefficient (Figure 4B), the anomalous diffusion parameter α^21^ (Figure 4C), and eGFP brightness as reflected by counts *per* second *per* molecule (Supplemental Figure S6), were determined. Of note, the fact that eGFP brightness in N2a cells did not change during hyperosmotic stress (Supplemental Figure S6), indicates that eGFP oligomerization did not occur to a considerable effect under hyperosmotic stress, and that eGFP remains largely monomeric and that observed changes in eGFP fluorescence lifetime are not due to phenomena such as homo-FRET.

**Figure 4.**
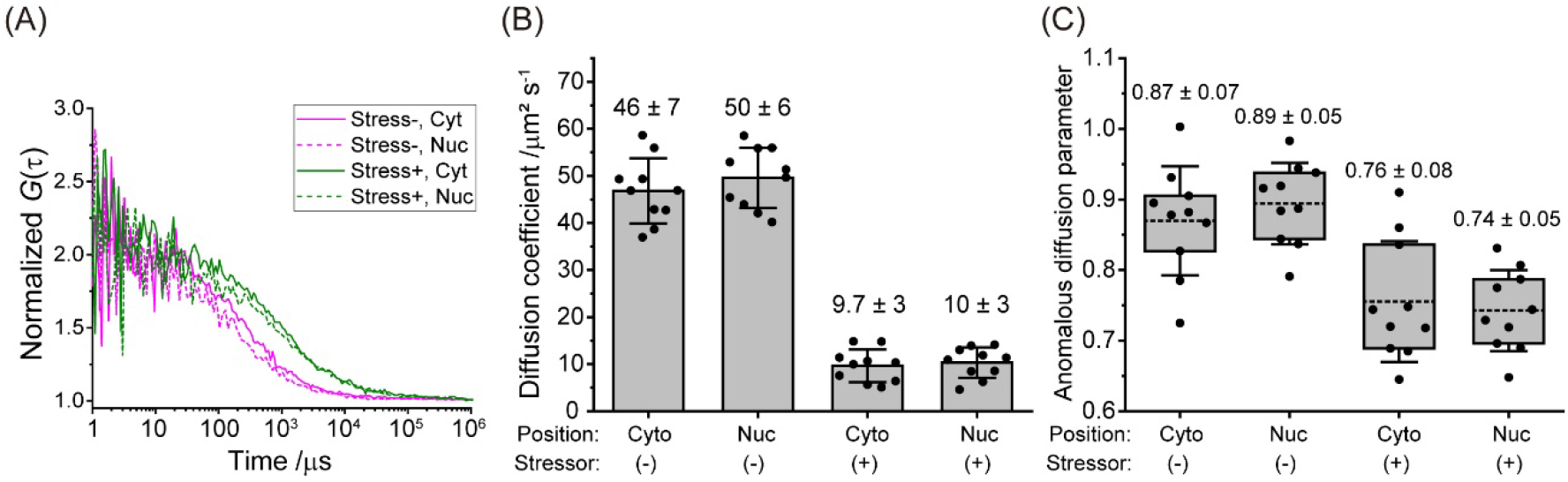
Hyperosmotic stress-induced changes in intracellular macromolecular crowding (MMC) slow down eGFP diffusion in live N2a cells and increase intracellular heterogeneity. (A) Autocorrelation curves normalized to the same amplitude, *G*(τ) = 1 at τ = 10 μs, reflecting differences in eGFP diffusion in live N2a cells before (Stress-) and after (Stress+) exposure to hyperosmotic stress, recorded in the cytoplasm (Cyt, solid line) and the nucleus (Nuc, dashed line).(B) Diffusion coefficient values determined by non-linear fitting of autocorrelation curves using Eq. 1 derived for anomalous three-dimensional diffusion of one-component with blinking due only to an equilibrium between dark and bright states. Bars and the numbers above them indicate mean ± SD (the number of analyzed cells, n = 10). (C) Box-and-whisker plot showing variations in the anomalous diffusion parameter (α) obtained by fitting Eq. 1 to autocorrelation curves shown in (A). The box indicates 25^th^ and 75^th^ percentile. The whisker indicates SD. The bold line shows mean value. The numbers above the bars indicate mean ± SD (the number of analyzed cells, n = 10).

The diffusion coefficient of eGFP monomers in N2a cells during hyperosmotic stress was significantly decreased, becoming 5-fold lower than that under normal, non-stress conditions. In agreement with previous observations^16^, differences between eGFP diffusion in the cytoplasm and the cell nucleus were not observed. Because viscosity is inversely proportional to the diffusion coefficient^8^, one could, based on the Stokes-Einstein equation, conclude for *in vitro* measurements that the viscosity increased 5-times. However, when it comes to measurements in live cells, one needs to be cautious in quantitative estimations because there are small contributions from numerous processes that on their own may be negligible but may exert a sizeable macroscopic net effect. Hence, we can only conclude that eGFP diffusion coefficient analysis suggests that hyperosmotic stress induces at most a 5-fold increase in intracellular viscosity *via* increased MMC. This interpretation is further corroborated by the following observations: (1) the anomalous diffusion parameter α, significantly decreased, as is expected when MMC and spatial heterogeneity in the cells increase^22^ (Figure 4B); (2) anomalous eGFP diffusion in BSA, which showed that α = (0.95 ± 0.05) in PBS, consistent with the expected largely free 3D diffusion of eGFP, while in 250 mg/ml BSA α = (0.86 ± 0.03), consistent with free diffusion hindrance due to increased viscosity due to MMC (Supplemental Figure S7); and (3) the 1.45 times increase in photon counts, which indicates that the cell volume decreased to ∽70 % of its initial size just after hyperosmotic stress (Supplemental Figure S2C), which is also in line with direct cell imaging data.

## Discussion

Our data show that both, τ_FL,eGFP_ and Y/C ratio of the GimRET sensor, decreased swiftly after exposing the N2a cells to hyperosmotic stress, and then gradually recovered, but remained still low after 3 hours of incubation in a hyperosmotic medium (Figures 1 and 2). This observation is in line with transmitted light microscopy imaging, which showed cell volume shrinkage consistent with dehydration shortly after exposing N2a cells to the hyperosmotic medium. It also showed that the cell volume did not recover throughout the observation time – in line with what was previously observed for other neuronal cells^23,24^. However, the half recovery time of eGFP fluorescence lifetime, τ_FL,eGFP_ and the Y/C ratio after stress medium addition were different, 9.7 s and 73.3 s, respectively, and eGFP lifetime changed faster than the GimRET Y/C ratio, reaching a minimum value a few seconds earlier (Figure 3). A possible explanation for this difference is that in GimRET conformational changes between the two linked fluorescent proteins affect FRET efficiency. The conformational rearrangement may be way slower than dynamic quenching of eGFP fluorescence, which may occur through a number of processes that increase radiation-less deactivation of eGFP, such as collisional quenching propelled by transiently increased concentration due to increased MMC and/or rapid reduction of cell volume. Since cells respond to hyperosmotic stress *via* protein phase separation^25^, eGFP monomers could “sense” such changes more readily than the dimeric GimRET since accessibility to secluded cell regions depends on protein size^26^.

Given that diffusion of non-reacting biomolecules is directly affected by the viscosity of the surrounding medium and that eGFP is not chemically reactive in cells, the significantly slower diffusion of monomeric eGFP is likely indicative of increased viscosity inside the cells primarily due to increased MMC triggered by the increase in biomolecular concentration^27^, rather than due to changes in eGFP conformation, activity and/or quinary interactions^28^, and the refractive index increment, caused by volume reduction due to exposure to hyperosmotic stress. Unfortunately, while we cannot provide a sole answer to the question to what extent different photophysical process, *e*.*g*. internal conversion or intersystem crossing to the triplet state, or oxidative photochemical reactions contribute to the observed decrease in τ_FL,eGFP_, we could show that the change in τ_FL,eGFP_ is reversible and that washing out the hyperosmotic medium is accompanied by τ_FL,eGFP_ return to its original value (Supplemental Figure S8).

Persistently high MMC in cells stimulates a direct apoptotic pathway^29^. For example, apoptosis signal-regulating kinase 3 (ASK3; also known as mitogen-activated protein kinase kinase kinase 15 [MAP3K15]) and its phosphorylation signaling play an important role in cell survival and osmotic stress response^30^. Since the MMC state in the cell affects kinase-substrate interaction^15^, high MMC during hyperosmotic stress regulates phosphorylation signaling activity associated with hyperosmotic stress responses. Although a quantitative measurement system and stable fluorescent proteins against such intracellular environmental changes are indispensable to characterize such change, concentration of several ions, and pH during dehydration easily change the photophysical property of fluorescent proteins, such as dynamic quenching, distorting the quantitative fluorescence measurement. An increase of intracellular MMC in *E*.*coli* during hyperosmotic stress was suggested using a FRET-based crowding sensor with a linker region that contains two α-helices and three random coils between a FRET pair: mCerulean3 and mCitrine, a chloride ion-insensitive YFP variant^31^; however, it has not been tested in mammalian cells. Furthermore, since we show that the recovery response of τ_FL,eGFP_ is faster than that of Y/C ratio of GimRET, these FRET-based crowding sensor may not show a rapid response due to the rate-determining step of structural change. We demonstrated that τ_FL,eGFP_ is sensitive to MMC change during hyperosmotic stress in mammalian cells, not affected by NaCl concentration and slight pH increase^18,32^. Unaltered eGFP brightness in N2a cells exposed to hyperosmotic stress (Supplemental Figure S6) and the high dissociation constant of eGFP (74 mM)^33^ suggest that significant eGFP homo-oligomerisation did not occur and support the interpretation that eGFP diffusion is reduced predominantly due to increased intercellular viscosity. Thus, FLIM using eGFP is a reliable method to measure MMC during hyperosmotic stress. Moreover, diffusion analysis by FCS is informative for determining intracellular viscosity. A somewhat shorter τ_FL,eGFP_ was observed in the nucleus than in the cytoplasm, while no difference was observed in diffusion state (Figures 1 and 4). We propose that MMC in the nucleus *via* chromatin condensation may be more susceptible to collisional quenching of eGFP. Therefore, a combination of both FLIM and FCS represents a more reliable method for determining MMC during hyperosmotic stress.

Accordingly, we demonstrate intracellular MMC and viscosity change during hyperosmotic stress using dynamic quenching of eGFP fluorescence. This strategy allows the quantitative analysis of various MMC states not only during hyperosmotic stress but also during other environmental changes in the cell.

## Methods

### Cell culture, transfection, and hyperosmotic medium

Murine neuroblastoma Neuro-2a (N2a) cells were obtained from American Type Culture Collection (ATCC; Manassas, VA) and maintained in DMEM (D5796, Sigma-Aldrich, St. Louis, MO) supplemented with 10% FBS (12676029, Thermo Fisher Scientific, Waltham, MA), 100 U/mL penicillin G (Sigma-Aldrich) and 0.1 mg/ml streptomycin (Sigma-Aldrich) as previously described^34^. One day before transfection 2.0 × 10^5^ N2a cells were transferred to a glass bottom dish (3910-035, IWAKI-AGC Technoglass, Shizuoka, Japan) for confocal imaging and FCS experiments or an 8-well chambered coverglass (155411, Thermo Fisher Scientific) for FLIM experiments. Plasmid DNAs were transfected using Lipofectamine 2000 (Thermo Fisher Scientific) according to the manufacturer’s protocol. After overnight incubation, the transfection medium was replaced with Opti-MEM I Reduced serum medium (Opti-MEM; 31985070, Thermo Fisher Scientific) for control experiments, which was thereafter replaced with Opti-MEM containing 290 mM NaCl for the hyperosmotic stress experiments.

### Plasmids

To express the enhanced green fluorescent protein carrying the A206K monomeric mutation (eGFP), peGFP-C1 and pTKbX-eGFP plasmids were used for FLIM and FCS, respectively^34^. The plasmid for the crowding-sensitive glycine-inserted mutant of YFP (YFP1G) fused with CFP (GimRET) sensor is described elsewhere^7^. The DNA fragment encoding the T-cell-restricted intracellular antigen-1 (TIA-1) was digested from pFRT-TO-eGFP-TIA1 (#106094, Addgene) using Bsp1407I (TaKaRa, Shiga, Japan) and inserted into mCherry-C1 (mCherry-TIA1).

### Multi-point fluorescence lifetime imaging microscopy (MP-FLIM)

The instrumental design and construction of the massively parallel imaging system without laser scanning is described previously^11,12^. Briefly, simultaneous excitation of fluorescent molecules across the sample is achieved by passing a single pulsed laser beam through a DOE, which transforms it into a rectangular illumination matrix that consists of 16×16 foci. Fluorescent signal from 256 foci in the illumination matrix is detected by a matrix detector, single-photon avalanche photodiodes (SPAD) camera. Fluorescence and bright field images are captured by the 18.0 megapixel digital single-lens reflex camera Canon EOS 600D (Canon Inc., Tokyo, Japan) to identify GFP localization and visualize cell morphology. FLIM data acquisition and analysis was carried out using the MPD SPC3 software (http://www.micro-photon-devices.com/Software) and the homebuilt MP-FLIM software^11,12^. Acquisition of a single FLIM curve lasted 3.7 s. τ_FL,eGFP_ was determined by fitting a single-exponential decay function to the experimental FLIM curves. To monitor the time course of FLIM changes, single FLIM curves were fitted and an average value from 9 spots in 5 cells is obtained for each time point (Figure 1D). To record data at steady-state conditions, that is before exposure to hyperosmotic stress and afterwards when a plateau value was reached, FLIM curves were acquired for 5 min and averaged to obtain low-noise FLIM curves at a single-pixel level and the average FLIM curve was fitted (Figure 1C).

### Fluorescence correlation spectroscopy (FCS)

To measure eGFP diffusion under normal physiology, FCS measurements were performed in N2a cells transiently expressing eGFP that were grown in the Opti-MEM medium (control) in a glass bottom dish. To measure eGFP diffusion under hyperosmotic stress, the Opti-MEM cell culture medium was replaced with the hyperosmotic stress medium, Opti-MEM with 290 mM NaCl. FCS measurements were performed using an LSM510 META ConfoCor3 system (Carl Zeiss, Jena, Germany) equipped with a C-Apochromat 40×/1.2NA W Korr. UV-VIS-IR water immersion objective, a multi-line Argon laser and Avalanche Photodiode (APD) detectors. eGFP was excited using the 488 nm laser line of the Argon laser. To remove the detector-derived afterpulsing noise, fluorescence was split using a half mirror (HF50/50), and detected, after passing through a 505-540 nm band-pass filter (BP505-540), using two APD detectors. Auto- and cross-correlation functions (ACFs and CCFs, respectively) were calculated for different lag times (τ) and plotted as a function of lag-time to derive corresponding auto- and cross-correlation curves. The cross-correlation curves were further analyzed by fitting a theoretical function for three-dimensional anomalous diffusion of one-component with blinking due only to an equilibrium between dark and bright states (Eq. 1) using the ZEN2012 (Carl Zeiss) software:

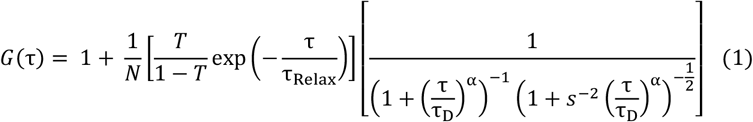

In Eq.1, *G*(τ) is the cross-correlation function; τ_D_ is the diffusion time of eGFP; α is the anomalous diffusion parameter^35^; *N* is the average number of eGFP molecules in the effective detection volume; *s* is the structure parameter representing the ratio of the radial (*w*_0_) and the axial (*z*_0_) 1/*e*^2^ radii of the effective volume; *T* is the exponential relaxation fraction; and τ_Relax_ is the relaxation time of eGFP blinking dynamics. In calibration experiments, the diffusion time (τ_Rh6G_) and structure parameter (*s*) were determined using 0.1 μM Rhodamine 6G (Rh6G) as a standard. The diffusion coefficient of eGFP was calculated using equation 2.

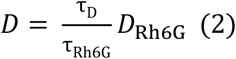

where *D* is the diffusion coefficient of eGFP; *D*_Rh6G_ is the diffusion coefficient of Rh6G (414 μm^2^/s); τ_D_ and τ_Rh6G_ are the diffusion times of eGFP and Rh6G, respectively. eGFP brightness, as reflected by counts *per* second *per* molecule, was calculated by dividing the mean count rate with the average number of molecules in the effective detection volume (*N*).

### Confocal fluorescence microscopy

Confocal images were acquired using a laser scanning microscope system LSM 510 META (Carl Zeiss) equipped with a C-Apochromat 40×/1.2NA W Korr. UV-VIS-IR water immersion objective, diode laser 405 nm, multi-line Argon and yellow He-Ne 594 nm laser, and photomultiplier tube (PMT) detectors. eGFP and mCherry were excited at 488 nm and 594 nm, respectively. Excitation and emission were separated using the main dichroic beam splitter HFT488/594. To spectrally separate eGFP and mCherry fluorescence, the secondary dichroic mirror NFT545 was used to split fluorescence emission at 545 nm. eGFP fluorescence was detected after passing through a 505-550 nm band-pass filter (BP505-550), and mCherry fluorescence was detected after passing through a 615 nm long pass filter (LP615). The pinhole was set to 71 μm for eGFP and 86 μm for mCherry. GimRET fluorescence was excited at 405 nm. Excitation and emission light were separated using the main dichroic beam splitter HFT405/514/594. Fluorescence was detected using a spectrometric multi-channel detector; the range 430-473 nm was collected for CFP and 558-655 nm for YFP. Pinhole was set to 300 μm. Acquired images were processed and analyzed using Fiji-ImageJ 1.52n. The Y/C ratio images were constructed by dividing in each pixel the background-subtracted fluorescence intensity of YFP with the background-subtracted fluorescence intensity of CFP and smoothed using the Gaussian-blur function in ImageJ.

## Acknowledgments

A.K. was supported by grants from the: Japan Society for Promotion of Science (JSPS) Grant-in-Aid for the Promotion of Joint International Research (Fostering Joint International Research) (16KK0156); JSPS Grant-in-Aid for Scientific Research (18K06201 and 22H04826); Japan Agency for Medical Research and Development (grants JP22gm6410028 and JP22ym0126814); Hoansha Foundation; Nakatani Foundation; Hagiwara Foundation; Canon Foundation; Japan Science and Technology Agency (JST) Competitive Funding Program for Adaptable and Seamless Technology Transfer Program through Target-driven R&D (A-STEP) (VP30318089120); Office for Developing Future Research Leaders (L-Station), Hokkaido University. M.K. was partially supported by JSPS Grant-in-Aid for Scientific Research (22H02578 and 22K19886). H.K. and M.O. were supported by a Hokkaido University Learning Satellite fellowship program for studying abroad. S.O. gratefully acknowledges a postdoctoral fellowship from the Nakatani Foundation for Advancement of Measuring Technologies in Biomedical Engineering and a travel grant from Yoshida Foundation for Science and Technology. V.V. gratefully acknowledges support from the Swedish Research Council (VR 2018-05337), the Olle Engkvist’s Foundation (Grant 199-0480) and the Magnus Bergvall Foundation (2021-04376). The funding agencies had no influence on the study design, methods, data collection, analyses or the manuscript writing.

## Author contribution

AK and MK designed the experiments and prepared the first draft of the manuscript. AK performed and analyzed confocal imaging using eGFP, GimRET, and mCherry-TIA1, in addition to preparation of all plasmids. HK and MO performed the FCS experiments and analyzed the data together with AK and MK. SO and HK performed the mpFLIM experiments and analyzed the data under the supervision of VV. The manuscript was written through contributions of all authors. All authors have given approval to the final version of the manuscript.

## Availability of materials and data

The datasets generated during and/or analysed during the current study are available in the Open science framework (OSF) repository, DOI: 10.17605/OSF.IO/FD2TA.

