## Supplemental Figures for "Increased intracellular crowding during hyperosmotic stress"

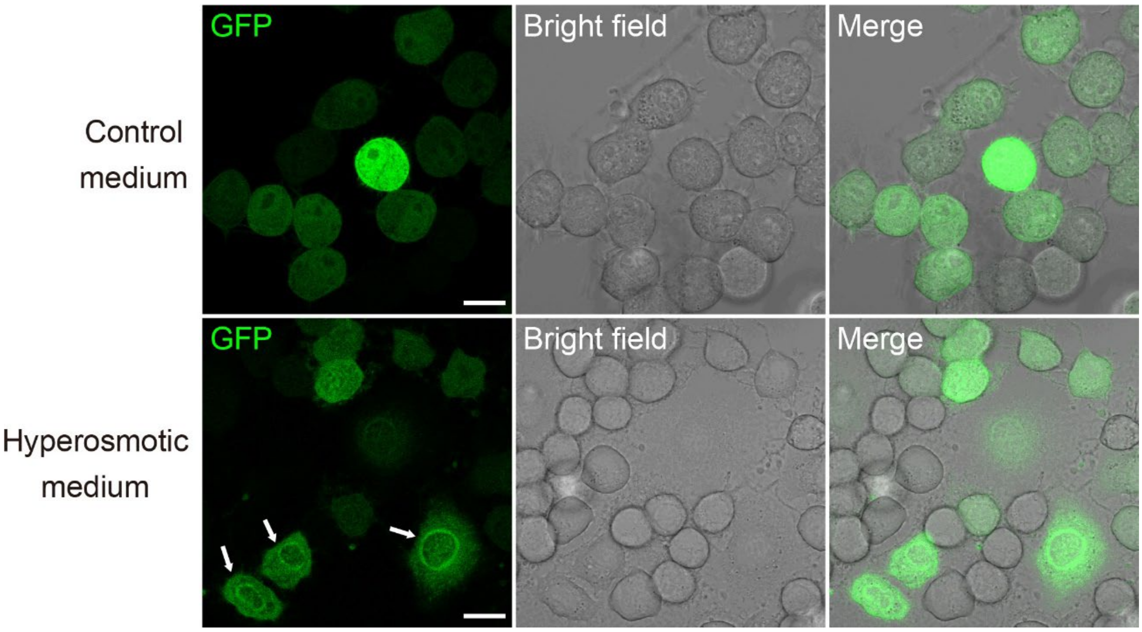


**Supplemental Figure S1. Redistribution of eGFP fluorescence in live N2a cells exposed to hyperosmotic stress.**

CLSM images showing eGFP fluorescence in live N2a cells before (upper row) and after (lower row) exposure to hyperosmotic stress. Note the intensely fluorescent rim around the nucleus in cells under hyperosmotic stress. Scale bar: 10 µm.

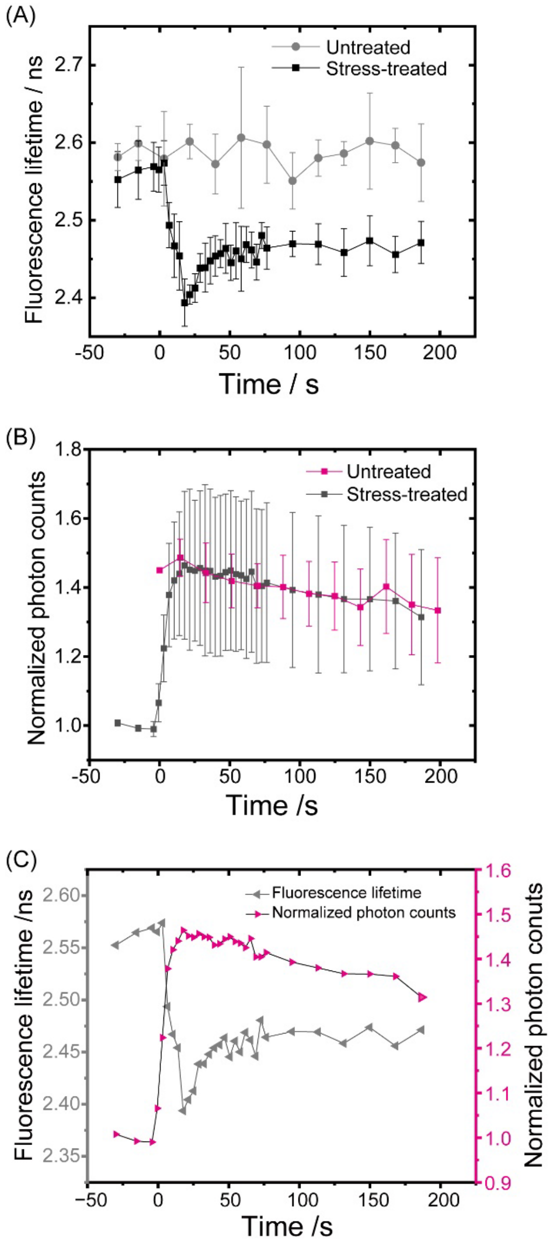


**Supplemental Figure S2. Changes over time in eGFP fluorescence lifetime and normalized photon counts measured in cells exposed and not exposed to hyperosmotic stress**

(A) eGFP fluorescence lifetime ($\tau_{FL,eGFP}$) in N2a cells exposed to hyperosmotic stress (black squares) and in not exposed N2a cells (grey dots). (B) Changes in fluorescence intensity, as reflected by the number of photon counts recorded during eGFP fluorescence lifetime measurements, normalized to the average value measured before exposure to hyperosmotic stress (black squares). In control cells (magenta squares), the photon counts are normalized to the value at $t = 0$ s, and a constant value of 0.42 is added to the values determined at each time point to shift upwards the curve and allow easier comparison with data acquired in cells exposed to hyperosmotic stress. Of note, the average photon counts recorded in the analysed live cells ranged 6.5 kHz – 65 kHz. (C) Change over time in fluorescence intensity recorded during $\tau_{FL,eGFP}$ measurement in live N2a cells exposed to hyperosmotic stress, as reflected by the normalized photon counts (magenta) and $\tau_{FL,eGFP}$ (gray). The hyperosmotic stress medium was added at $t = 0$ s. Dots and bars show mean ± SD (the number of analysed spots from 5 cells, n = 9). The graph showing changes over time in $\tau_{FL,eGFP}$ measured in cells exposed to hyperosmotic stress that is shown in A & C is the same as in Figure 1D.

(A)

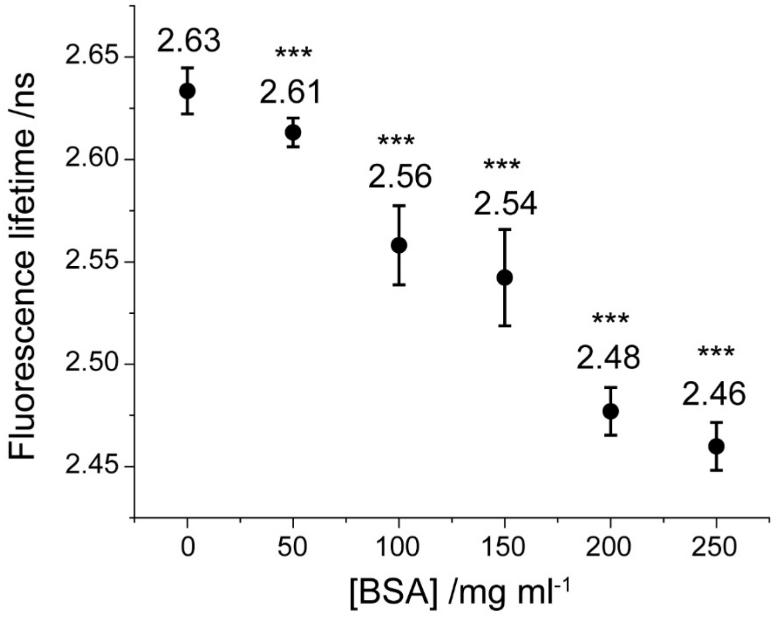


(B)

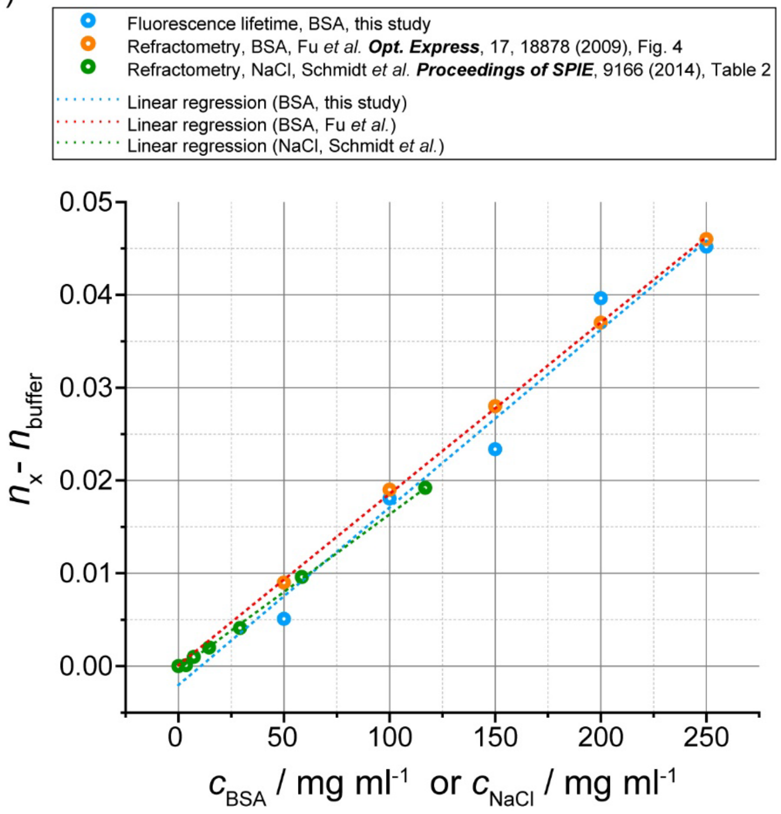


**Supplemental Figure S3. BSA-induced macromolecular crowding (MMC) in buffer solution causes BSA concentration-dependent changes in eGFP fluorescence lifetime**

(A) Fluorescence lifetime of monomeric eGFP as a function of BSA concentration in phosphate buffered saline (PBS), pH 7.4, at room temperature (21 °C). Dots and bars show mean ± SD, respectively (number of spots analyzed, n = 256). Statistical significance is determined with regard to measurements in PBS alone (no BSA added) using Student's *t* test: $***p < 0.001$. (B) Refractive index increment as a function of BSA concentration ($c_{\mathrm{BSA}}$) calculated from refractive index measurements reported by Fu D. *et al.*, ***Opt. Express*** (2009), Figure 4, DOI: 10.1364/OE.17.018878 (orange dots); calculated from eGFP fluorescence lifetime shown in (A) using the relationship $n_x = n_{\mathrm{buffer}} \cdot \sqrt{\tau_{\mathrm{FL,eGFP,buffer}}/\tau_{x\mathrm{FL,eGFP},x}}$ that is derived based on the Strickler–Berg equation (cyan dots); and refractive index increment as a function of NaCl concentration ($c_{\mathrm{NaCl}}$) calculated based on measured refractive index by Schmidt *et al.* ***Proceedings of SPIE*** (2014), Table 2, DOI:10.1117/12.2062389 (green dots).

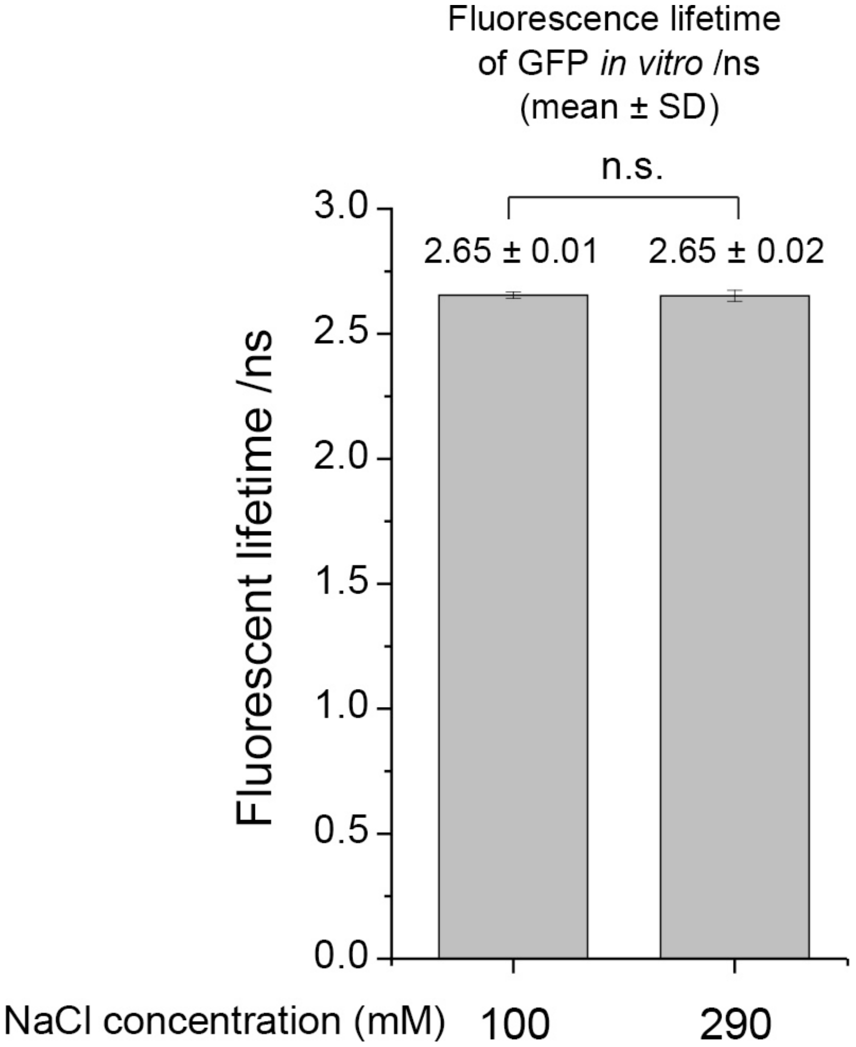


**Supplemental Figure S4. eGFP fluorescence lifetime in solution is not affected by NaCl**

Fluorescence lifetime of monomeric eGFP in phosphate buffered saline (PBS), pH 7.4, at room temperature (21 °C), is the same in 100 mM (physiological level) and 290 mM (hyperosmotic stress level) NaCl solution. The numbers on the bar graph show mean ± SD (the number of spots analyzed, n = 256). Statistical significance analysis, performed using the Student's *t* test, showed that statistical significance was not observed (n.s., $p > 0.05$).

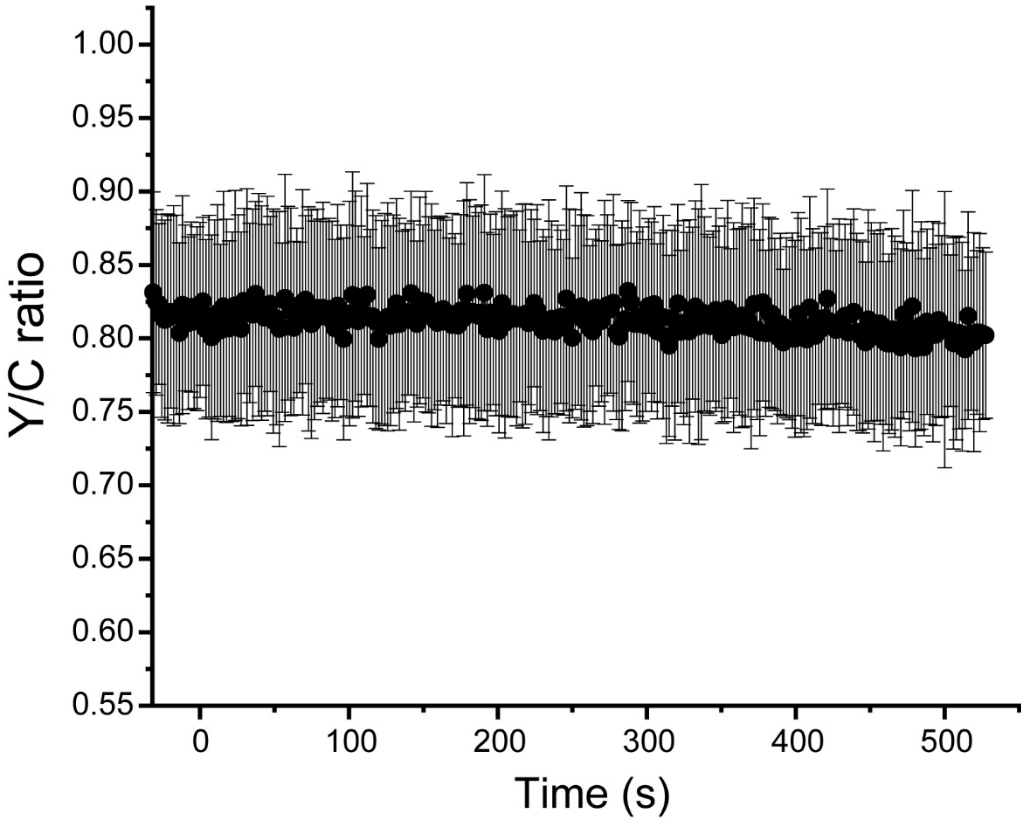


**Supplemental Figure S5. The Y/C ratio in GimRET-expressing N2a cells that were not exposed to hyperosmotic stress is not changing over time**

The Y/C ratio was determined by dividing the background-corrected fluorescence intensity in the YFP channel by the background-corrected fluorescence intensity in the CFP channel. Dots and bars indicate mean ± SD (the number of analyzed cells, n = 15).

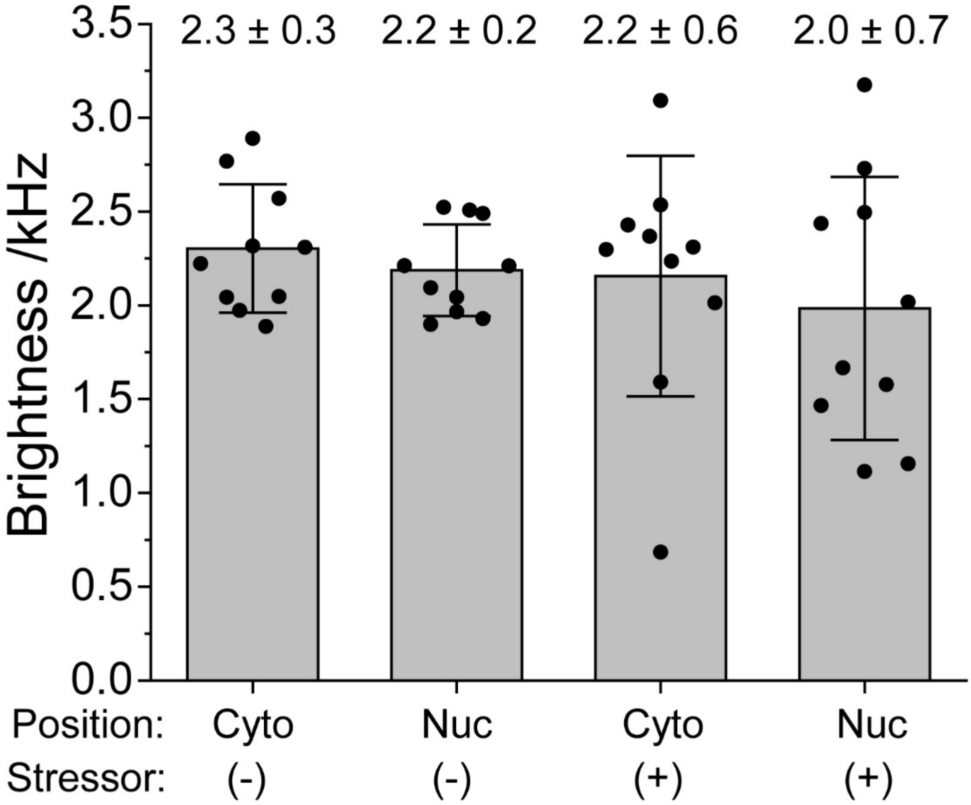


**Supplemental Figure S6. eGFP brightness in live N2a cells is not affected by hyperosmotic stress**
eGFP brightness, as reflected by counts *per* second *per* molecule measured by FCS. The numbers on the bar graph show mean ± SD (the number of spots analyzed, n = 10). Statistical significance analysis, performed using the Student's t test, showed that statistical significance was not observed ($p > 0.05$).

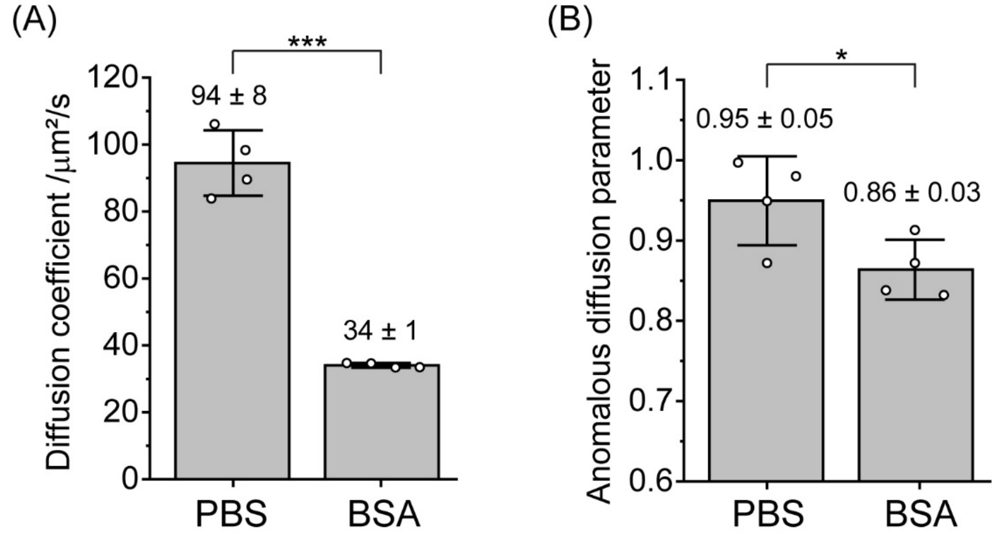


**Supplemental Figure S7. eGFP diffusion coefficient and anomalous diffusion parameter measured in PBS and 250 mg/ml BSA solution**

**(A)** Diffusion coefficient and **(B)** anomalous diffusion parameter (α) of purified eGFP monomers in phosphate buffer saline (PBS, pH 7.4), and 250 mg/ml bovine serum albumin (BSA) were measured at room temperature (21 °C) using FCS. Diffusion coefficient values determined by non-linear fitting of autocorrelation curves using Eq. 1 derived for anomalous three-dimenional diffusion of one-component with blinking due only to one equilibrium between dark and bright states. Bars and the numbers above them indicate mean ± SD (n = 4). Statistical significance analysis, performed using the Student's *t* test, showed statistical significance (*, $p < 0.05$; ***, $p < 0.001$).

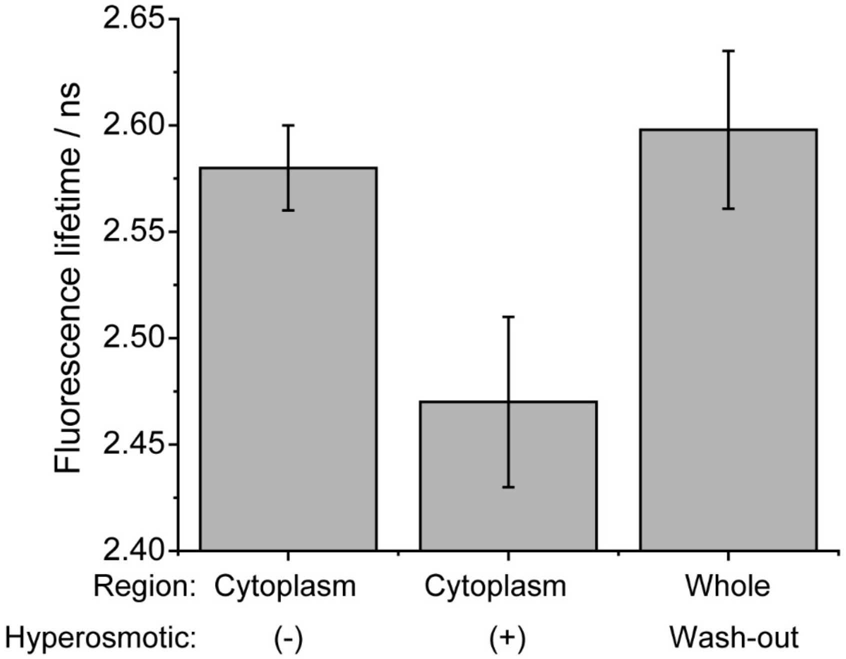


**Supplemental Figure S8. eGFP fluorescence lifetime in live N2a cells reversibly changes under hyperosmotic stress.**

eGFP fluorescence lifetime in live N2a cells before hyperosmotic stress, 2.58 ± 0.02; after hyperosmotic stress, 2.47 ± 0.04; and after washing out the hyperosmotic medium, 2.60 ± 0.04 (the number of cells analyzed, n = 10).
